# Simple penalties on maximum likelihood estimates of genetic parameters to reduce sampling variation

**DOI:** 10.1101/034447

**Authors:** Karin Meyer

**Author notes:** AGBU is a joint venture of NSW Department of Primary Industries and the University of New England.

## Abstract

Multivariate estimates of genetic parameters are subject to substantial sampling variation, especially for smaller data sets and more than a few traits. A simple modification of standard, maximum likelihood procedures for multivariate analyses to estimate genetic covariances is described, which can improve estimates by substantially reducing their sampling variances. This is achieved maximizing the likelihood subject to a penalty. Borrowing from Bayesian principles, we propose a mild, default penalty – derived assuming a Beta distribution of scale-free functions of the covariance components to be estimated – rather than laboriously attempting to determine the stringency of penalization from the data. An extensive simulation study is presented demonstrating that such penalties can yield very worthwhile reductions in loss, i.e. the difference from population values, for a wide range of scenarios and without distorting estimates of phenotypic covariances. Moreover, mild default penalties tend not to increase loss in difficult cases and, on average, achieve reductions in loss of similar magnitude than computationally demanding schemes to optimize the degree of penalization. Pertinent details required for the adaptation of standard algorithms to locate the maximum of the likelihood function are outlined.

## INTRODUCTION

Estimation of genetic parameters, i.e. partitioning of phenotypic variation into its causal components, is one of the fundamental tasks in quantitative genetics. For multiple characteristics of interest, this involves estimation of covariance matrices due to genetic, residual and possibly other random effects. It is well known that such estimates can be subject to substantial sampling variation. This holds especially for analyses comprising more than a few traits, as the number of parameters to be estimated generally increases quadratically with the number of traits considered, unless the covariance matrices of interest have a special structure and can be modelled more parsimoniously. Indeed, a sobering but realistic view is that “Few datasets, whether from livestock, laboratory or natural populations, are of sufficient size to obtain useful estimates of many genetic parameters” (Hill, 2010; p.75). This emphasizes not only the importance of appropriate data, but also implies that a judicious choice of methodology for estimation – which makes the most of limited and precious records available – is paramount.

A measure of the quality of an estimator is its ‘loss’, i.e. the deviation of the estimate from the true value. This is an aggregate of bias and sampling variation. We speak of improving an estimator if we can modify it so that the expected loss is lessened. In most cases, this involves reducing sampling variance at the expense of some bias – if the additional bias is small and the reduction in variance sufficiently large, the loss is reduced. In statistical parlance ‘regularization’ refers to the use of some kind of additional information in an analysis. This is often used to solve ill-posed problems or to prevent over-fitting through some form of penalty for model complexity, see Bickel and Li (2006) for a review. There has been longstanding interest, dating back to Stein (1975) and earlier (James and Stein, 1961) in regularized estimation of covariance matrices to reduce their ‘loss’. Recently, as estimation of higher dimensional matrices is becoming more ubiquitous, there has been a resurgence in interest (e.g. Bickel and Levina, 2008; Warton, 2008; Witten and Tibshirani, 2009; Ye and Wang, 2009; Rothman *et al*., 2010; Fisher and Sun, 2011; Ledoit and Wolf, 2012; Deng and Tsui, 2013; Won *et al*., 2013).

### Improving estimates of genetic parameter

As emphasized above, quantitative genetic analyses require at least two covariance matrices to be estimated, namely due to additive genetic and residual effects. The partitioning of the total variation into its components creates substantial sampling correlations between them and tends to exacerbate the effects of sampling variation inherent in estimation of covariance matrices. However, most studies on regularization of multivariate analyses considered a single covariance matrix only and the literature considering regularized estimates of more than one covariance matrix is sparse. In a classic paper, Hayes and Hill (1981) proposed to modify estimates of the genetic covariance matrix (**Σ***_G_*) by shrinking the canonical eigenvalues of **Σ***_G_* and the phenotypic covariance matrix (**Σ***_P_*) towards their mean, a procedure they described as ‘bending’ the estimate of **Σ***_G_* towards that of **Σ***_P_*. The underlying rationale was that **Σ***_P_*, the sum of all the causal components, is typically estimated much more accurately than any of its components, so that bending would ‘borrow strength’ from the estimate of **Σ***_P_*, while shrinking estimated eigenvalues towards their mean would counteract their known, systematic overdispersion. The authors demonstrated by simulation that use of ‘bent’ estimates in constructing selection indices could increase the achieved response to selection markedly, as these were closer to the population values than unmodified estimates and thus provided more appropriate estimates of index weights. However, no clear guidelines to determine the optimal amount of shrinkage to use were available and ‘bending’ was thus primarily used only to modify non-positive definite estimates of covariance matrices, and all but forgotten when methods which allowed estimates to be constrained to the parameter space became common procedures.

Modern analyses to estimate genetic analyses are generally carried out fitting a mixed model and using restricted maximum likelihood (REML) or Bayesian methodology. The Bayesian framework directly offers the opportunity for regularization through the choice of appropriate priors. Yet, this is rarely exploited for this purpose and ‘flat’ or minimally informative priors are often used instead (Thompson *et al*., 2005). In a maximum likelihood context, estimates can be regularized by imposing a penalty on the likelihood function aimed at reducing their sampling variance. This provides a direct link to Bayesian estimation: for a given prior distribution of the parameters of interest or functions thereof, an appropriate penalty can be obtained as a multiple of minus the logarithmic value of the probability density function. For instance, shrinkage of eigenvalues towards their mean through a quadratic penalty on the likelihood is equivalent to assuming a Normal distribution of the eigenvalues while the assumption of a double exponential prior distribution results in a LASSO type penalty (Huang *et al*., 2006).

Meyer and Kirkpatrick (2010) demonstrated that a REML equivalent to ‘bending’ can be obtained by imposing a penalty proportional to the variance of the canonical eigenvalues on the likelihood, derived assuming a Normal distribution of the eigenvalues with common mean, and showed that this can yield substantial reductions in loss for estimates of both **Σ***_G_* and **Σ***_E_*, the residual covariance matrix. Subsequent simulations (Meyer *et al*., 2011; Meyer, 2011) examined the scope for penalties based on different functions of the parameters to be estimated and prior distributions for them and found them to be similarly effective, depending on the population values for the covariance matrices to be estimated.

A central component of the success of regularized estimation is the choice of how much to penalize. A common practice is to scale the penalty by a so-called ‘tuning factor’ to regulate stringency of penalization. Various studies (again for a single covariance matrix; see above for references) demonstrated that this can be estimated reasonably well from the data at hand using cross-validation techniques. Adopting these suggestions for genetic analyses and using *k*–fold cross-validation (for *k* = 3 or 5), Meyer (2011) estimated the appropriate tuning factor as that which maximised the average, unpenalized likelihood in the validation sets. However, this procedure was laborious and afflicted by problems in locating the maximum of a fairly flat likelihood surface for analyses involving many traits and not so large data sets. These technical difficulties all but prevented practical applications. Moreover, it was generally less successful than reported for studies considering a single covariance matrix. This led to the suggestion of imposing a mild penalty, determining the tuning factor as the largest value which did not cause a decrease in the (unpenalized) likelihood equivalent to a significant change in a single parameter. This pragmatic approach yielded reductions in loss which were generally of comparable magnitude to those achieved using cross-validation (Meyer, 2011). However, it still required multiple analyses and thus considerably increased computational demands compared to standard, unpenalized estimation.

### Simple penalties

In the Bayesian framework, the influence of the prior and thus the amount of regularization is generally specified through the so-called hyperparameters of the prior distribution which determine its shape, scale or location. This suggests that an alternative, tuning factor-free formulation for the penalty on the likelihood can be obtained by expressing it in terms of the distribution-specific (hyper)parameters which are not dependent on the covariance matrices. For instance, when assuming a Normal prior for canonical eigenvalues, the regulating parameter is the variance of the Normal distribution, with more shrinkage induced the lower its value. This may lend itself to applications employing default values for these parameters. Furthermore, such formulation may facilitate direct estimation of the regulating parameter, denoted henceforth as *ν*, simultaneously with the covariance components to be estimated (de los Campos 2013; pers. comm.). In contrast, in a setting involving a tuning factor the penalized likelihood is, by definition, highest for a factor of zero (i.e. no penalty), and thus does not provide this opportunity.

This paper examines the scope for REML estimation imposing penalties regulated by choosing the parameters of the selected prior distribution. The focus is on penalties involving scale-free functions of covariance components which fall into defined intervals and may thus be better suited to a choice of default regulating parameters than functions which are not. We begin with the description of suitable penalties together with a brief review of pertinent literature and outline the adaptation of standard REML algorithms. This is followed by a large-scale simulation study showing that the penalties proposed can yield substantial reductions in loss of estimates for a wide range of population parameters. We conclude with a discussion and recommendations on selection of default parameters for routine use in multivariate analyses.

## PENALIZED MAXIMUM LIKELIHOOD ESTIMATION

Consider a simple mixed, linear model for *q* traits with covariance matrices **Σ***_G_* and **Σ***_E_* due to additive genetic and residual effects, respectively, to be estimated. Let 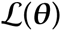 denote the log likelihood in a standard, unpenalized maximum likelihood (or REML) analysis and *θ* the vector of parameters, comprised of the distinct elements of **Σ***_G_* and **Σ***_E_* or equivalent. The penalized likelihood is then (Green, 1987)

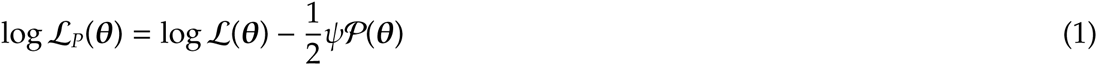

with the penalty 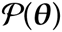 a non-negative function of the parameters to be estimated and *ψ* the so-called tuning factor which modulates the strength of penalization (the factor of ½ is used for algebraic consistency and could be omitted). In the following, we assume *ψ* = 1 throughout and regulate the amount of penalization instead via the parameters of the distribution from which the penalty is derived.

### Functions to be penalized

We consider two types of scale-free functions of the covariance matrices to be estimated as the basis for regularization.

**Canonical eigenvalues:** Following Hayes and Hill (1981), the first comprises the canonical eigenvalues of **Σ***_G_* and **Σ***_P_* = **Σ***_G_* + **Σ***_E_*.

Multivariate theory shows that for two symmetric, positive definite matrices of the same size there is a transformation which yields **TT**′ = **Σ***_P_* and **T**Λ**T**′ = **Σ***_G_*, with Λ the diagonal matrix of canonical eigenvalues with elements *λ_i_* (Anderson, 1984). This can be thought of as transforming the traits considered to new variables which are uncorrelated and have phenotypic variance of unity, i.e. the canonical eigenvalues are equal to heritabilities on the new scale and fall in the interval [0, 1] (Hayes and Hill, 1980). It is well known that estimates of eigenvalues of covariance matrices are systematically biased – the largest values are overestimated and the smallest are underestimated – while their mean is expected to be estimated correctly (Lawley, 1956). Moreover, a major proportion of the sampling variation of covariance matrices can be attributed to this over-dispersion of eigenvalues (Ledoit and Wolf, 2004). Hence there have been various suggestions to modify the eigenvalues of sample covariance matrices in some way to reduce the loss in estimates; see Meyer and Kirkpatrick (2010) for a more detailed review.

**Correlations:** The second type of functions comprises correlations between traits, in particular genetic correlations. A number of Bayesian approaches to the estimation of covariance matrices decompose the problem into variances (or standard deviations) and correlations with separate priors, thus alleviating the inflexibility of the widely used conjugate prior given by an Inverse Wishart distribution (Barnard *et al*., 2000; Daniels and Kass, 2001; Zhang *et al*., 2006; Daniels and Pourahmadi, 2009; Hsu *et al*., 2012; Bouriga and Féron, 2013; Gaskins *et al*., 2014). On the whole, however, few suitable families of prior density functions for correlation matrices have been considered and estimation using Monte Carlo sampling schemes has been hampered by difficulties arising from the constraints of positive definiteness and unit diagonals.

Most statistical literature concerned with Bayesian or penalized estimation of correlation matrices considered shrinkage towards an identity matrix, i.e. shrinkage of individual correlations towards zero, though other, simple correlation structures have been proposed (Schäfer and Strimmer, 2005). As outlined above, the motivation for ‘bending’ (Hayes and Hill, 1981) included the desire to ‘borrow’ strength from the estimate of the phenotypic covariance matrix. Similar arguments may support shrinkage of the genetic towards the phenotypic correlation matrix. This dovetails with what has become known as ‘Cheverud’s conjecture’: Reviewing a large body of literature Cheverud (1988), found that estimates of genetic correlations were generally close to their phenotypic counterparts, and thus proposed that phenotypic values should be substituted when genetic correlations could not be estimated. Subsequent studies reported similar findings for a range of traits in laboratory species, plants and animals (e.g. Roff, 1995; Waitt and Levin, 1998; Koots *et al*., 1994).

### Partial correlations

Often a reparameterisation can transform a constrained matrix problem to an unconstrained one. For instance, it is common practice in REML estimation of covariance matrices to estimate the elements of their Cholesky factors, coupled with a logarithmic transformation of the diagonal elements, to remove constraints on the parameter space (Meyer and Smith, 1996). Pinheiro and Bates (1996) examined various transformations for covariance matrices and their impact on convergence behaviour of maximum likelihood analyses, and corresponding forms for correlation matrices have been described (Rapisarda *et al*., 2007). Joe (2006) proposed a reparameterisation of correlations to partial correlations which vary independently over the interval [−1, 1], with a one-to-one transformation between partial and standard correlations. This implies, that it is possible to sample a random correlation matrix which is positive definite by sampling individual partial correlations. Daniels and Pourahmadi (2009) referred to these quantities as partial auto-correlations (PAC), interpreting them as correlations between traits *i* and *j* conditional on the intervening traits, *i* + 1 to *j* − 1.

Consider a correlation matrix **R** of size *q* × *q* with elements *ρ_ij_* (for *i* ≠ *j*) and *ρ_ii_* = 1. As **R** is symmetric, let *i* < *j*. For *j* = *i* + 1, the PAC are equal to the standard correlations, *π_i_*,*_i_*_+1_ = *ρ_i_*,*_i_*_+1_, as there are no intervening variables. For *j* > *i* + 1, partition the submatrix of **R** comprised of rows and columns *i* to *j* as

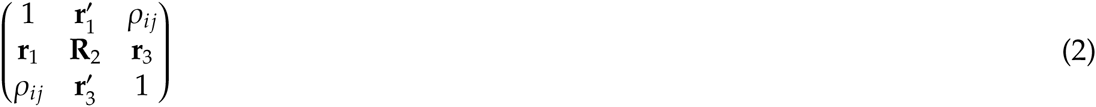

with **r**_1_ and **r**_3_ vectors of length *j* − *i* − 1, with elements *ρ_ik_* and *ρ_jk_*, respectively, and **R**_2_ the corresponding matrix with elements *ρ_kl_* for *k*, *l* = *i* + 1,…, *j* − 1. This gives PAC

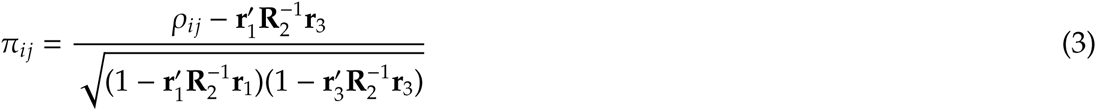

and the reverse transformation is

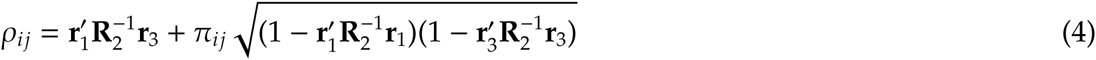

(Joe, 2006).

### Penalties

We derive penalties on canonical eigenvalues and correlations or partial auto-correlations assuming independent Beta distributions as priors.

**Beta distribution:** The Beta distribution is a continuous probability function that is widely used in Bayesian analyses and encompasses functions with many different shapes, determined by two parameters, *α* and *β*. While the standard Beta distribution is defined for the interval [0, 1], it is readily extended to a different interval. The probability density function for a variable *x* ∈ [*a*, *b*] following a Beta distribution is of the form

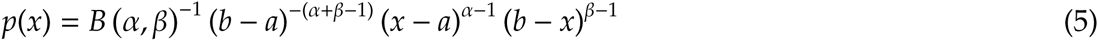

(Johnson *et al*., 1995; Chapter 25) with *B*(*α*, *β*) = Γ (*α*) Γ(*β*)/Γ(*α* + *β*) and *B* (·) and Γ (·) denoting the Beta and Gamma function, respectively.

When employing a Beta prior in Bayesian estimation, the sum of the shape parameters, *ν* = *α* + *β*, is commonly interpreted as the effective sample size (ESS) of the prior (Morita *et al*., 2008). It follows that we can specify the parameters of a Beta distribution with mode *m* as a function of the ESS (*ν* ≥ 2) as

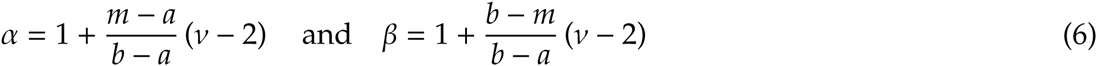

For *ν* > 2, this yields a unimodal distribution, and for *m* = (*b* − *a*)/2 the distribution is symmetric, with *α* = *β*. For given *m*, this provides a mechanism to regulate the strength of penalization through a single, intuitive parameter, the ESS *ν*.

Figure 1 shows the probability density of a variable with a standard Beta distribution on the interval [0, 1] with mode of 0.3 together with the resulting penalty, for three values of ESS. For *ν* = 2, the density function would be a uniform distribution, depicted by a horizontal line at height of 1, resulting in no penalty. With increasing values of *ν*, the distribution becomes more and more peaked and the penalty on values close to the extremes of the range becomes more and more severe. Conversely, in spite of substantial differences in point mass around the mode, penalty values in proximity of the mode differ relatively little for different values of *ν*. While 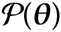 in (1) was considered non-negative, penalty values close to the mode can be negative – this does not affect suitability of the penalty and can be overcome by adding a suitable constant.

**Figure 1:**
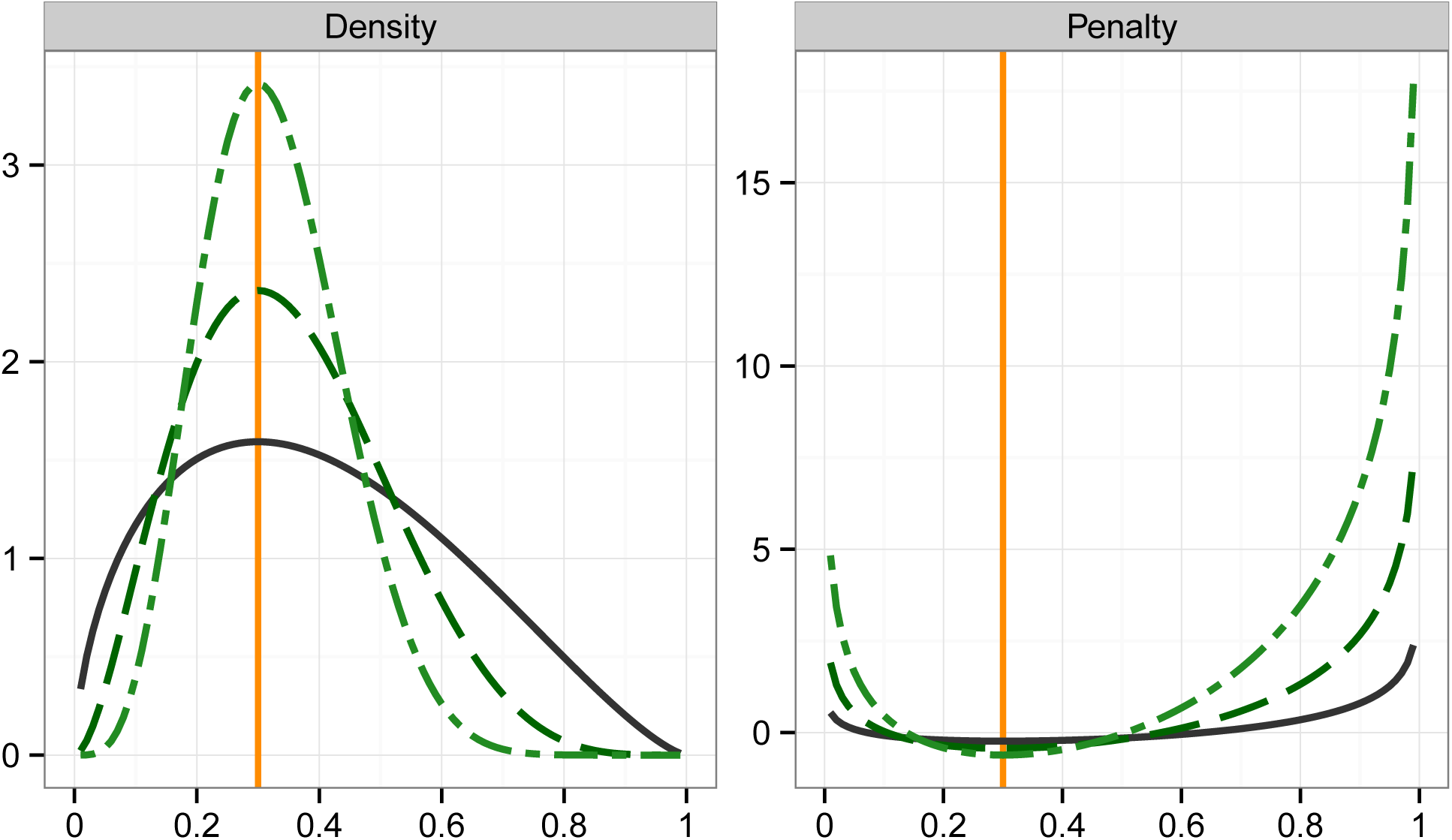
Probability densities (left) and corresponding penalties (including a factor of ½; right) for a variable with Beta distribution on [0, 1] with mode of *m* = 0.3 for effective sample sizes of *ν* = 4(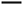), *ν* = 8(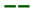)and *ν* = 16(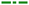).

**Penalty on canonical eigenvalues:** Canonical eigenvalues fall in the interval [0, 1]. For *q* traits, there are likely to be *q* different values *λ_i_* and attempts to determine a mode may be futile. Hence we propose to substitute the mean canonical eigenvalue, 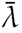, for the mode. Taking minus logarithmic values of (5) and assuming the same mode and ESS for all *q* eigenvalues then gives penalty

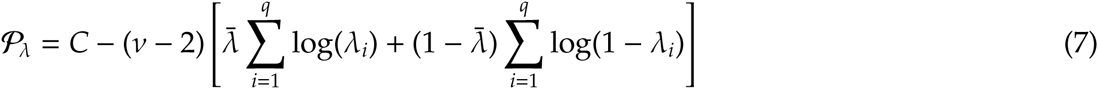

with 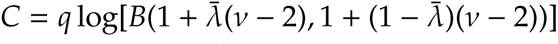. This formulation results in shrinkage of all eigenvalues towards 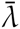.

**Penalty on correlations:** For (standard) correlations and PAC, we assume independent shifted Beta distributions on [−1, 1].

Both Joe (2006) and Daniels and Pourahmadi (2009) considered such Beta priors for PAC with *α* = *β*, i.e. shrinkage of all *π_ij_* towards zero. We generalize this by allowing for different shrinkage targets *τ_ij_* – and thus different shape parameters *α_ij_* and *β_ij_* – for individual values *π_ij_*. This gives penalty

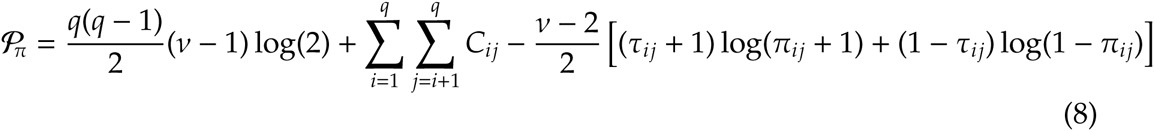

with *C_ij_* = log[*B*(1 + (*τ_ij_* + 1)(*ν* − 2)/2, 1 + (1 − *τ_ij_*)(*ν* − 2)/2)]. Again, this assumes equal ESS for all values, but could of course readily be expanded to allow for different values *ν_ij_* for different PAC. For all *τ_ij_* = 0, (8) reduces to

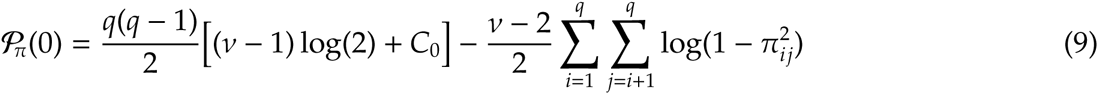

with *C*_0_ = log[*B*(1 + (*ν* − 2)/2, 1 + (*ν* − 2)/2)]. Corresponding penalties on standard correlations, 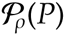 and 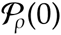, are obtained by substituting *ρ_ij_* for *π_ij_* in (8) and (9).

Daniels and Pourahmadi (2009) considered several Bayesian priors for correlation matrices formulated via PAC, suggesting uniform distributions for individual *π_ij_*, i.e. *π_ij_* ~ Beta(1, 1). In addition, they showed that the equivalent to the joint uniform prior for **R** proposed by Barnard *et al*. (2000), *p*(**R**) ∝ 1, is obtained by assuming Beta priors for PAC with shape parameters depending on the number of intervening variables, i.e. *α_i_*,*_i_*_+_*_k_* = *β_i_*,*_i_*_+_*_k_* = 1 + (*q* − 1 − *k*)/2. Similarly, priors proportional to higher powers of the determinant of **R**, *p*(**R**) ∝ |**R**|*^t^*^−1^, are obtained for *α_i_*,*_i_*_+_*_k_* = *t* + (*q* − 1 − *k*)/2 (Daniels and Pourahmadi, 2009). Gaskins *et al*. (2014) extended this framework to PAC based priors with more aggressive shrinkage towards zero for higher lags, suitable to encourage sparsity in estimated correlation matrices for longitudinal data.

### Maximizing the penalized likelihood

REML estimation in quantitative genetics usually relies on algorithms exploiting derivatives of the log likelihood function to locate its maximum. In particular, the so-called average information algorithm (Gilmour *et al*., 1995) is widely used due to its relative computational ease, good convergence properties and implementation in several REML software packages. It can be described as Newton(-Raphson) type algorithm where the Hessian matrix is approximated by the average of observed and expected information. To adapt the standard, unpenalized algorithm for penalized estimation we need to adjust first and second derivatives of 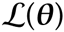 for derivatives of the penalties with respect of the parameters, *θ_k_*, to be estimated. These differ if we choose fixed values to determine the modes of the assumed Beta priors (e.g. *τ_ij_* = 0 or the mean *λ_i_* from a preliminary, unpenalized analysis) and or employ penalties which derive these from the parameter estimates.

Consider the penalty on canonical eigenvalues. If 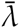 is estimated from the data, its derivatives are non-zero. This gives

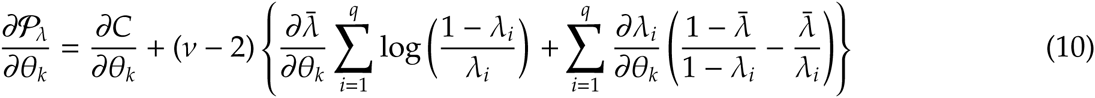

and

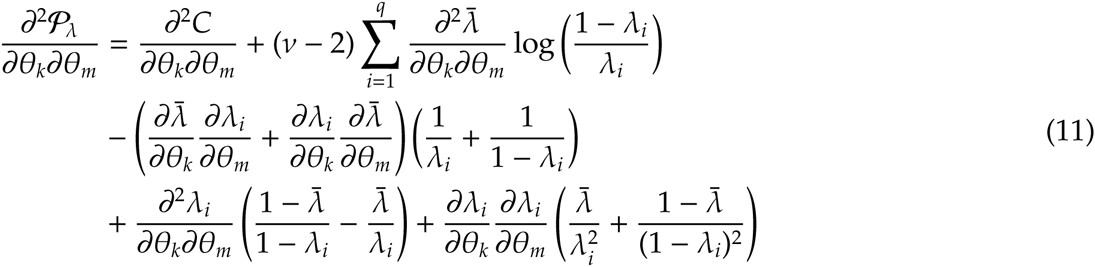

Derivatives of *C* involve the Digamma and Trigamma functions, e.g.

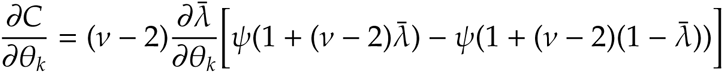

with *ψ* the Digamma function. Derivatives of *λ_i_* required in (10) and (11) are easiest to evaluate if the analysis is parameterised to the canonical eigenvalues and the elements of the corresponding transformation matrix **T** (see Meyer and Kirkpatrick, 2010), so that 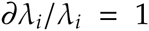 that 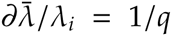 and 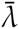 and all other derivatives of 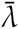 and 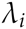 are zero. A possible approximation is to ignore contributions of derivatives of 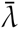, arguing that the mean eigenvalue is expected to be unaffected by sampling over-dispersion and thus should change little.

Analogous arguments hold for penalties involving correlations. This gives

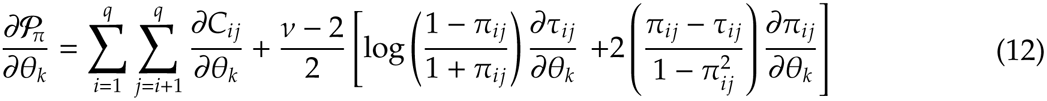

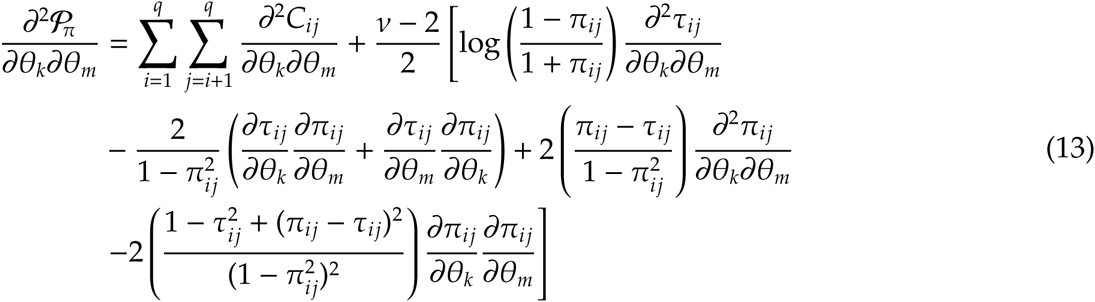

with obvious simplifications if shrinkage targets are fixed or treated as such, so that derivatives of *τ_ij_* are zero. As shown in the appendix, derivatives of correlations and PACs are readily calculated from the derivatives of covariance components for any of the parameterizations commonly utilised in (unpenalized) REML algorithms for variance component estimation.

## SIMULATION STUDY

A large scale simulation study, considering a wide range of population parameters, was carried out to examine the efficacy of the penalties proposed above.

### Set-up

Data were sampled from multivariate normal distributions for *q* = 9 traits, assuming a balanced paternal half-sib design comprised of *s* unrelated sire families with 10 progeny each. Sample sizes considered were *s* = 100, 400 and 1000, with records for all traits for each of the progeny but no records for sires.

Population values for genetic and residual variance components were generated by combining 13 sets of heritabilities with six different types of correlation structures to generate 78 cases. Details are summarized in the appendix and population canonical eigenvalues for all sets are shown in Supplement S1. To assess the potential for detrimental effects of penalized estimation, values were chosen deliberately to generate both cases which approximately matched the priors assumed in deriving the penalties and cases where this was clearly not the case. The latter included scenarios where population canonical eigenvalues were widely spread and in multiple clusters and cases where genetic and phenotypic correlations were highly dissimilar. A total of 500 samples per case and sample size were obtained.

### Analyses

REML estimates of **Σ***_G_* and **Σ***_E_* for each sample were obtained without penalization and imposing different penalties, namely a penalty on canonical eigenvalues 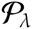, as given in (7) and a penalties on partial auto-correlations shrinking all values towards zero (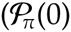, see (9)) and with shrinkage targets for each value equal to the corresponding phenotypic counterpart (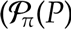, see (8)). For the latter two, penalties on genetic PAC only and both genetic and residual values were examined. In addition, penalties on standard correlations, shrinking towards zero 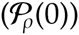 or phenotypic values 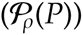 were considered for comparison.

Analyses were carried out considering fixed values for the effective sample size, ranging from *ν* = 2.5 to 24. For penalties on both genetic and residual correlations/PAC, either the same value was used for both or the ESS for residual correlation/PAC was fixed at *ν* = 8. In addition, direct estimation of a suitable ESS for each replicate was attempted. As shown in (1), penalties were subtracted from the standard log likelihood incorporating a factor of 1/2.

The model of analysis was a simple animal model, fitting means for each trait as the only fixed effects. A Method of Scoring algorithm together with simple derivative-free search steps was used to locate the maximum of the (penalized) likelihood function. To facilitate easy computation of derivatives of 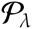, this was done using a parameterisation to the elements of the canonical decomposition (see Meyer and Kirkpatrick, 2010), restraining estimates of *λ_i_* to the interval of [0.0001, 0.9999]. For penalties on correlations, a parameterisation to the elements of the Cholesky factors of **Σ***_G_* and **Σ***_E_* was used, constraining estimates of diagonal elements to a minimum of 0.0001.

Direct estimation of *ν* was performed by evaluating points on the profile likelihood for *ν* (i.e. maximizing 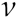 with respect to the covariance components to be estimated for selected, fixed values of *ν*), combined with quadratic approximation steps of the profile to locate its maximum using Powell’s (2006) Fortran subroutine <m>NEWUOA</m>. To avoid numerical problem, estimates of *ν* were constrained to the interval [2.01, 50].

### Summary statistics

For each sample and analysis, the quality of estimates was evaluated through their entropy loss (James and Stein, 1961)

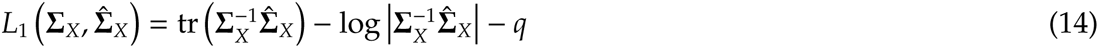

for *X* = *G*, *E* and *P*, with **Σ***_X_* denoting the matrix of population values for genetic, residual and phenotypic covariances, and 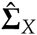 the corresponding estimate. As suggested by Lin and Perlman (1985), the percentage reduction in average loss (PRIAL) was used as the criterion to summarize the effects of penalization

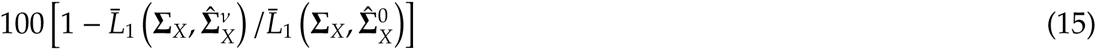

where 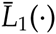 denotes the entropy loss averaged over replicates, and 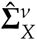 and 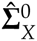 represent the penalized and corresponding unpenalized REML estimate of **Σ***_X_*, respectively.

In addition, the average reduction in log 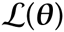 due to penalization, 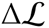, was calculated as the mean difference across replicates between the unpenalized likelihood for estimates 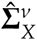 and the corresponding value for estimates 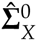.

## RESULTS

Distributions of PRIAL across the 78 sets of population values together with the corresponding change in likelihood are shown in Figure 2 for two sample sizes, with penalties 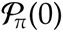 and 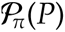 applied to genetic PAC only. Distributions given are trimmed, i.e. the range shown reflects the minimum and maximum values observed. Selected mean and minimum values are also reported in Table 1. More detailed results, including distributions for penalties 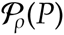 and 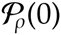 and values for individual cases, are reported in Supplement S1.

**Figure 2:**
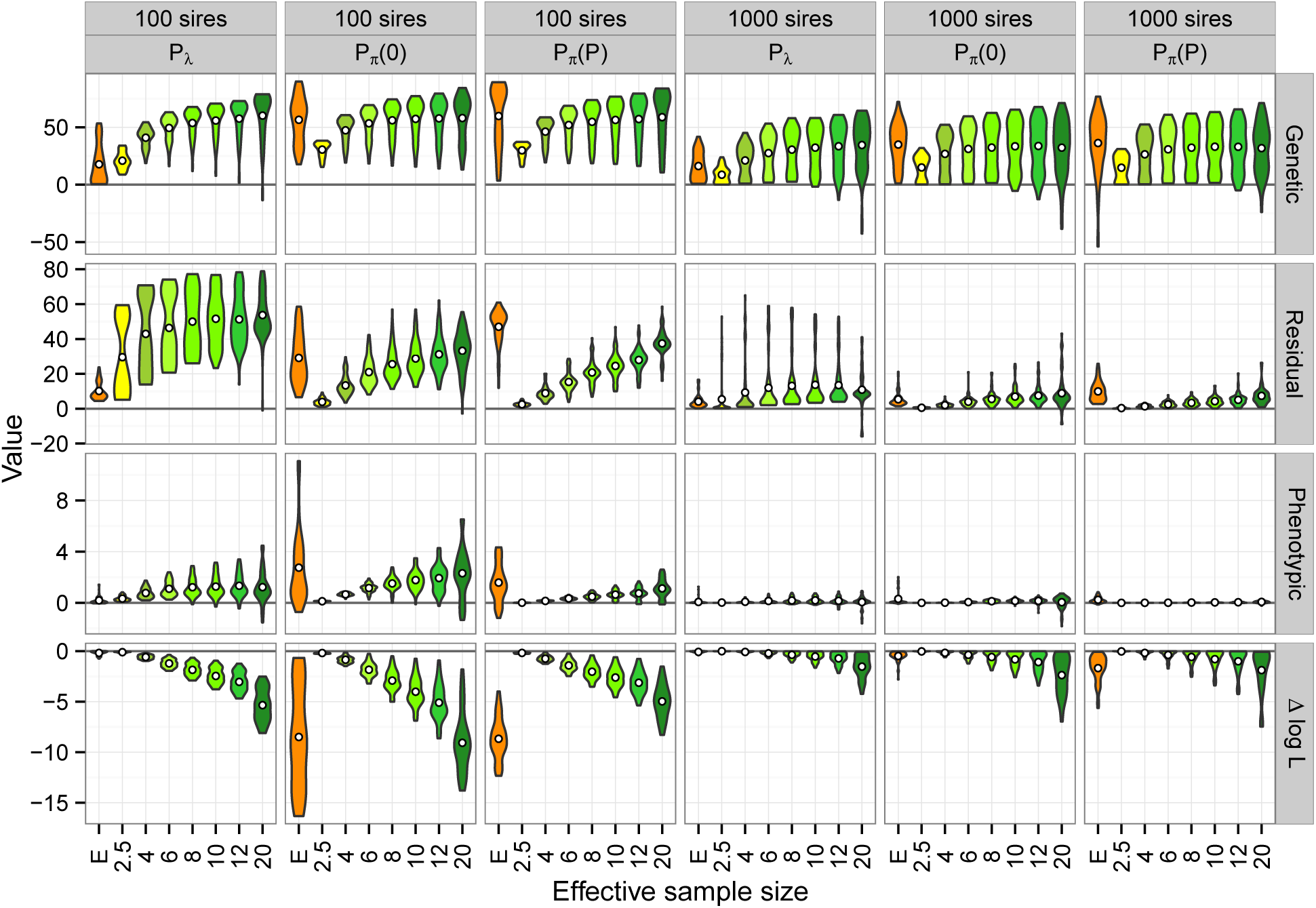
Distribution of percentage reduction in average loss for estimates of genetic, residual and phenotypic covariance matrices, together with corresponding change in log likelihood 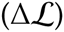 for penalties on canonical eigenvalues 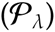 and genetic, partial auto-correlations, shrinking towards zero 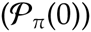 or phenotypic values 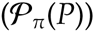. Central circles give mean values. Numeric values on the x-axis are fixed, effective sample sizes while “E” denotes the use of a value estimated from the data for each replicate.

**Table 1:** Selected mean and minimum values for percentage reduction in average loss for estimates of genetic (**Σ***_G_*), residual (**Σ***_E_*) and phenotypic (**Σ***_P_*) covariance matrices together with mean change in unpenalised log likelihood from the maximum 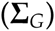 for penalties on canonical eigenvalues 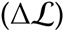 and genetic correlations, shrinking partial auto-correlations towards zero 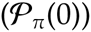 or phenotypic values 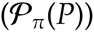 and standard correlations towards phenotypic values 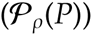.

| Penalty | $\nu^a$ | 100 sires | | | | 400 sires | | | | 1000 sires | | | |
| --- | --- | --- | --- | --- | --- | --- | --- | --- | --- | --- | --- | --- | --- |
| | | $\Sigma_G$ | $\Sigma_E$ | $\Sigma_P$ | $\Delta\mathcal{L}$ | $\Sigma_G$ | $\Sigma_E$ | $\Sigma_P$ | $\Delta\mathcal{L}$ | $\Sigma_G$ | $\Sigma_E$ | $\Sigma_P$ | $\Delta\mathcal{L}$ |
| Mean values |  |  |  |  |  |  |  |  |  |  |  |  |  |
| $\mathcal{P}_\lambda$ | 4 | 41 | 43 | 1 | -0.6 | 31 | 19 | 0 | -0.2 | 21 | 9 | 0 | -0.1 |
|  | 6 | 49 | 46 | 1 | -1.2 | 38 | 22 | 0 | -0.5 | 27 | 12 | 0 | -0.2 |
|  | 8 | 54 | 50 | 1 | -1.9 | 42 | 24 | 0 | -0.7 | 30 | 13 | 0 | -0.4 |
|  | 12 | 58 | 51 | 1 | -3.0 | 45 | 25 | 0 | -1.4 | 33 | 13 | 0 | -0.7 |
|  | E | 18 | 10 | 0 | -0.2 | 22 | 7 | 0 | -0.1 | 16 | 4 | 0 | -0.1 |
| $\mathcal{P}_\pi(0)$ | 4 | 47 | 13 | 1 | -0.9 | 37 | 5 | 0 | -0.3 | 27 | 2 | 0 | -0.2 |
|  | 6 | 53 | 21 | 1 | -1.8 | 43 | 9 | 0 | -0.7 | 31 | 4 | 0 | -0.4 |
|  | 8 | 56 | 26 | 2 | -2.9 | 44 | 12 | 0 | -1.1 | 32 | 6 | 0 | -0.6 |
|  | 12 | 58 | 31 | 2 | -5.1 | 45 | 16 | 1 | -2.1 | 34 | 8 | 0 | -1.1 |
|  | E | 56 | 29 | 3 | -8.5 | 46 | 12 | 1 | -1.9 | 35 | 5 | 0 | -0.5 |
| $\mathcal{P}_\pi(P)$ | 4 | 46 | 9 | 0 | -0.7 | 37 | 3 | 0 | -0.3 | 26 | 1 | 0 | -0.2 |
|  | 6 | 52 | 15 | 0 | -1.4 | 42 | 6 | 0 | -0.7 | 31 | 2 | 0 | -0.4 |
|  | 8 | 55 | 21 | 0 | -2.0 | 44 | 8 | 0 | -1.0 | 32 | 3 | 0 | -0.6 |
|  | 12 | 57 | 28 | 1 | -3.1 | 45 | 11 | 0 | -1.6 | 33 | 5 | 0 | -1.0 |
|  | E | 60 | 47 | 2 | -8.7 | 50 | 22 | 1 | -3.5 | 36 | 10 | 0 | -1.7 |
| $\mathcal{P}_\rho(P)$ | 4 | 8 | 11 | 0 | -0.3 | 13 | 5 | 0 | -0.2 | 11 | 2 | 0 | -0.1 |
|  | 8 | 20 | 21 | 0 | -1.3 | 24 | 10 | 0 | -0.9 | 20 | 5 | 0 | -0.6 |
|  | 12 | 28 | 26 | 0 | -2.1 | 29 | 12 | 0 | -1.7 | 23 | 6 | 0 | -1.2 |
|  | E | 42 | 38 | 0 | -4.9 | 39 | 19 | 0 | -2.9 | 29 | 6 | 0 | -1.8 |
| Minimum values |  |  |  |  |  |  |  |  |  |  |  |  |  |
| $\mathcal{P}_\lambda$ | 8 | 12 | 26 | 0 | -2.9 | 5 | 7 | 0 | -1.7 | 2 | 2 | 0 | -1.1 |
|  | 12 | 1 | 13 | 0 | -4.7 | -11 | 11 | -1 | -3.0 | -14 | 4 | -1 | -2.2 |
|  | E | 0 | 4 | 0 | -0.7 | 1 | 1 | 0 | -0.6 | 1 | 0 | 0 | -0.4 |
| $\mathcal{P}_\pi(0)$ | 8 | 16 | 11 | 0 | -5.0 | 2 | 2 | 0 | -3.0 | 1 | 0 | 0 | -1.9 |
|  | 12 | 14 | 11 | 0 | -8.7 | -7 | 2 | -1 | -5.4 | -13 | 0 | -1 | -3.4 |
|  | E | 18 | 7 | -1 | -16.4 | 1 | 3 | -1 | -11.6 | 1 | 1 | 0 | -2.8 |
| $\mathcal{P}_\pi(P)$ | 8 | 18 | 7 | 0 | -3.5 | 4 | 1 | 0 | -2.9 | 1 | 0 | 0 | -2.6 |
|  | 12 | 16 | 12 | 0 | -5.4 | -1 | 2 | 0 | -4.7 | -5 | 0 | 0 | -4.3 |
|  | E | 3 | 12 | -1 | -12.4 | -34 | 7 | 0 | -8.8 | -54 | 3 | 0 | -5.6 |
| $\mathcal{P}_\rho(P)$ | 8 | -27 | 7 | -1 | -3.6 | -7 | 1 | 0 | -3.9 | 1 | 0 | 0 | -3.4 |
|  | 12 | -36 | 10 | -1 | -5.2 | -15 | -5 | -1 | -7.2 | -3 | -9 | -1 | -6.2 |
|  | E | -33 | 6 | -2 | -8.6 | -21 | 1 | -1 | -7.3 | -27 | -13 | -2 | -5.3 |

<sup>a</sup>Effective sample size of the Beta distribution; numerical values are ‘fixed’ values used while E denotes values estimated from the data for each replicate, with means of 3.1, 5.9 and 8.5 (for 100, 400 and 1000 sires, respectively) for $\mathcal{P}_\lambda$ , 31.2, 16.0, 13.9 for $\mathcal{P}_\pi(0)$ and 45.4, 36.4 and 34.8 for $\mathcal{P}_\pi(P)$ and 34.0, 33.6. and 32.2 for $\mathcal{P}_\rho(P)$

### Genetic covariances

Overall, for fixed ESS there were surprisingly little differences between penalties 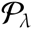, 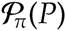 and 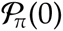, in mean PRIAL values achieved, especially for estimates of the genetic covariance matrix. Correlations between PRIAL for 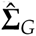 from the three different penalties ranged from 0.9 to unity, suggesting similar modes of action.

However, corresponding values for penalties on standard correlations were consistently lower and, more importantly, some minimum values were negative, even for small values of the ESS. As demonstrated in more detail in Supplement S1, this was due to marked mismatches between population values and shrinkage targets (*a.k.a*. priors) for one of the six correlation structures, which was deliberately included to test the robustness of the penalties proposed. Transformation to partial auto-correlations produced a better match and thus yielded penalties markedly less likely to have detrimental effects. While easier to interpret than PAC, penalties on standard correlations should just be used cautiously. In the following, we consider only penalties on canonical eigenvalues and PAC.

Even for small values of *ν* there were worthwhile reduction in loss for estimates of **Σ***_G_*, especially for the smallest sample (*s* = 100). Means increased with increasing stringency of penalization along with an increasing spread in results for individual cases, especially for the largest sample size. This pattern was due to the range of population values for genetic parameters used.

Moreover, for small samples or low ESS, it did not appear to be all that important whether the priors on which the penalties were based approximately matched population values or not: ‘any’ penalty proved beneficial, i.e resulted in positive PRIAL for 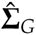 (except penalties on standard correlations for some cases). For more stringent penalization, however, there was little improvement (or even adverse effects) for the cases where there was a clear mismatch. For instance, for 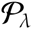 for *ν* = 24 and *s* = 100 sires, there were two cases with negative PRIAL for 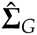. Both of these had a cluster of high and low population values for *λ_i_* so that the assumption of a unimodal distribution invoked in deriving 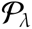 was inappropriate and led to sufficient over-shrinkage to be detrimental. On the whole, however, unfavourable effects of penalization were few and restricted to the most extreme cases considered.

Paradoxically, PRIAL for 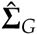 were also low for cases where heritabilities were approximately the same and genetic and phenotypic correlations were similar, so that canonical eigenvalues differed little from their mean (see S1). This could be attributed to the shape of the penalty function, as illustrated in Figure 1, resulting in little penalization for values close to the mode. In other words, these were cases were the prior did not quite match the population values: while the assumption of a common mean for canonical eigenvalues clearly held, that of a distribution on the interval [0, 1] did not. This can be rectified by specifying a more appropriate interval. As the unpenalized estimates of *λ_i_* are expected to be overdispersed, their extremes may provide a suitable range to be used. Additional simulations (not shown) for 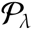 replaced values of *a* = 0 and *b* = 1 used to derive 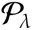 in 7) with 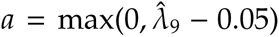 and 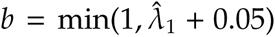 for each replicate, where 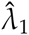 and 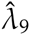 represented the largest and smallest canonical eigenvalue estimate from a preliminary, unpenalized analysis, respectively. This increased PRIAL for both 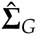 and 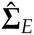 substantially for these cases. However, as the proportion of such cases overall was low (see S1), overall results were little affected.

### Residual covariances

Some differences between penalties were apparent for **Σ***_E_*. 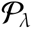 involved terms log(1 − *λ_i_*) (see (7)), i.e. the canonical eigenvalues of **Σ***_E_* and **Σ***_P_*. Hence, 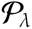 yielded substantial PRIAL for estimates of both **Σ***_G_* and **Σ***_E_*, especially for the smaller samples where sampling variances and losses were high. Conversely, applying penalties on genetic PAC resulted in some, but lower improvements in **Σ***_E_* (except for large *ν*), but only as a by-product due to negative sampling correlations between **Σ***_G_* and **Σ***_E_*. As shown in Table 2, imposing a corresponding penalty on residual PACs in addition could increase the PRIAL in estimates of **Σ***_E_* markedly without reduction in the PRIAL for **Σ***_G_*, provided the ESS chosen corresponded to a relatively mild degree of penalization. Shrinking towards phenotypic PAC yielded somewhat less spread in PRIAL for **Σ***_E_* than shrinking towards zero, accompanied by smaller changes in log 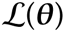.

**Table 2:**
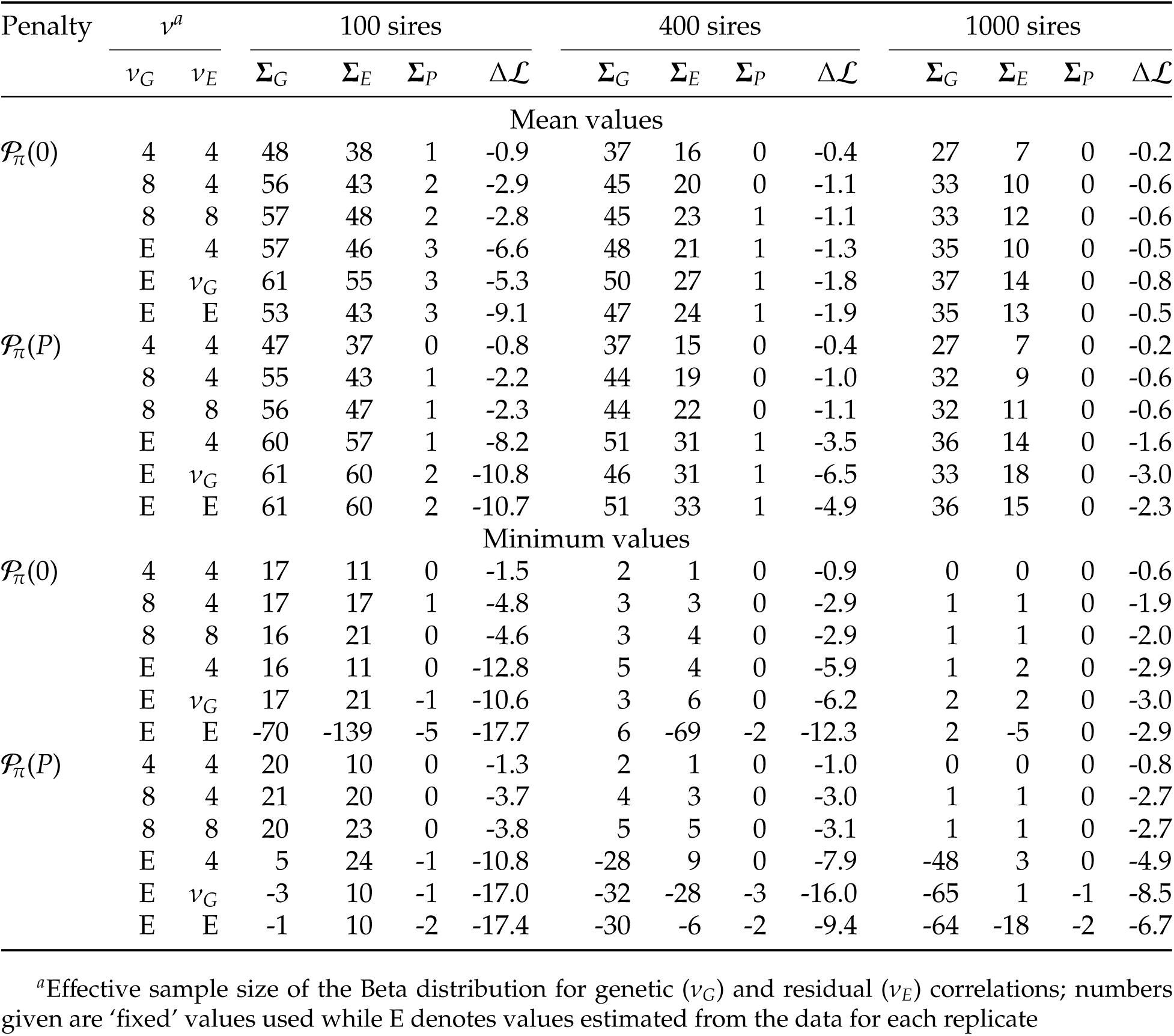
Selected mean and minimum values for percentage reduction in average loss for estimates of genetic (**Σ***_G_*), residual (**Σ***_E_*) and phenotypic (**Σ***_P_*) covariance matrices together with mean change in unpenalised log likelihood from the maximum 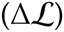 for penalties on both genetic and residual partial auto-correlations, shrinking towards zero 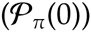 or phenotypic values 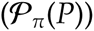

### Phenotypic covariances

We argued above that imposing penalties based on the estimate of **Σ***_P_* would allow us to ‘borrow strength’ because it typically is estimated more precisely than any of its components. Doing so, we would hope to have little – and certainly no detrimental – effect on the estimates of **Σ***_P_* as, loosely speaking, we would expect penalized estimation to redress, to some extent at least, any distortion in partitioning of **Σ***_P_* due to sampling correlations. As demonstrated in Figure 2, this was generally the case for fixed values of the ESS less than about *ν* = 10 or 12, with negative PRIAL for estimates of **Σ***_P_* for higher values flagging over-penalization where population values for genetic parameters did not sufficiently match the assumptions on which the penalties were based.

### Canonical eigenvalues

Figure 3 shows the distribution of the largest and smallest canonical eigenvalues, contrasting population values with mean estimates from unpenalized and penalized analyses for a medium sample size and a fixed ESS of *ν* = 8. Results clearly illustrate the upwards bias in estimates of the largest and downwards bias of the smallest eigenvalues. As expected, imposing penalty 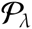 reduced the mean of the largest and increased the mean of the smallest eigenvalues, with some over-shrinkage, especially of the largest eigenvalue, evident. In contrast, for the small value of *ν* = 8 chosen, the distribution of the largest values from penalized and unpenalized analyses differed little for penalty 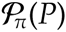, i.e. penalizing genetic PAC did not affect the leading canonical eigenvalues markedly, acting predominately on the smaller values. For more stringent penalties, however, some shrinkage of the leading eigenvalues due to penalties 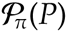 and 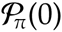 was evident; detailed results for selected cases are given in Supplement S1.

**Figure 3:**
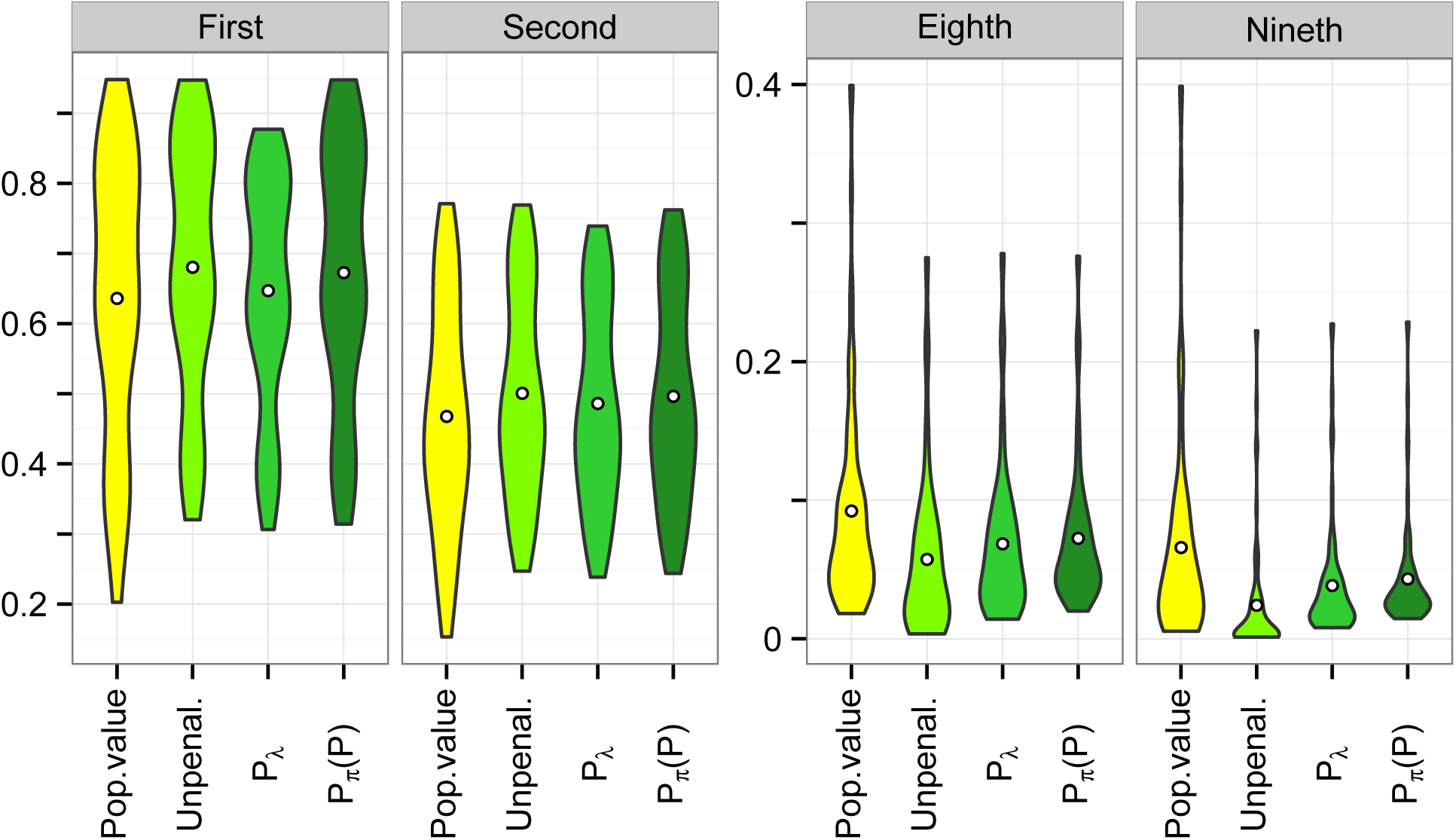
Distribution of mean estimates of selected canonical eigenvalues comparing results from unpenalized analyses and analyses imposing penalties on canonical eigenvalues (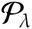) and genetic, partial auto-correlations, shrinking towards phenotypic values (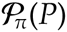), with population values in the simulation. Both penalties employ a fixed, effective sample size of *ν* = 8; results shown are for samples with 400 sire families. Central circles give mean values across the 78 cases considered.

### Estimating ESS

Overall, attempts to estimate the appropriate value of *ν* from the data were not all that successful. For 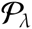, numerous cases yielded an estimate of *ν* close to the lower end of the range allowed, i.e. virtually no penalty. Conversely, for 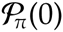 and 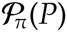 a substantial number of cases resulted in estimates of *ν* close to the upper bound allowed. This increased PRIAL (compared to fixed values for *ν*) for cases which approximately matched the priors but caused reduced or negative PRIAL and substantial changes in 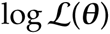 otherwise. A possible explanation was that the penalized likelihood and thus the estimate of *ν*, might be dominated by **Σ***_E_*. However, as shown in Table 2, neither estimating a value for **Σ***_G_*(*ν_G_*) while fixing the ESS for **Σ***_E_*(*ν_E_*) or estimating a value for both (either separately or jointly, *ν_G_* = *ν_E_*) improved results greatly. Moreover, it yielded more cases for which penalization resulted in substantial, negative PRIAL, especially for 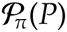. Repeating selected analyses using a sequential grid search to determine optimal values of *ν* gave essentially the same results, i.e. results could not be attributed to inadequacies in the quadratic approximation procedure.

## DISCUSSION

Sampling variation is the bane of multivariate analyses in quantitative genetics. While nothing can replace large numbers of observations with informative data and appropriate relationship structure, we often need to obtain reasonably trustworthy estimates of genetic parameters from relatively small data sets. This holds especially for data from natural populations but is also relevant for new or expensive to measure traits in livestock improvement or plant breeding schemes. We have shown that regularized estimation in a maximum likelihood framework through penalization of the likelihood function can provide ‘better’ estimates of covariance components, i.e. estimates that are closer to the population values than those from standard, unpenalized analyses. This is achieved through penalties targeted at reducing sampling variation.

Moreover, we have demonstrated that it is feasible to choose default values for the strength of penalization which yield worthwhile reductions in loss for a wide range of scenarios and are robust, i.e. are unlikely to result in penalties with detrimental effects, and are technically simple. While such tuning-free approach may not yield a maximum reduction in loss, it appears to achieve a substantial proportion thereof in most cases with modest changes in the likelihood compared to the maximum of 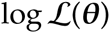 (without penalization). In contrast to attempts to estimate a tuning factor, it does not require multiple additional analyses to be carried out, and effects of penalization on computational requirements are thus mostly unimportant. In addition, we can again make the link to Bayesian estimation, where the idea of mildly or weakly informative priors has been gaining popularity. Discussing priors for variance components in hierarchical models, Gelman (2006) advocated a half-t or half-Cauchy prior with large scale parameter. Huang and Wand (2013) extended this to prior distributions for covariance matrices which resulted in half-t priors for standard deviations and marginal densities of correlations *ρ* proportional to a power of (1 − *ρ*^2^). Chung *et al*. (2015) proposed a prior distribution for covariance matrices proportional to a Wishart distribution with diagonal scale matrix and low degrees of belief and used this to obtain a penalty on the likelihood function which ensured non-degenerate estimates of variance components.

Results suggest that penalties on canonical eigenvalues or PAC assuming a Beta prior with a conservative choice of ESS of *ν* = 4 to 10 will result in substantial improvements in estimates of genetic covariance components for many cases, with little chance of detrimental effects for cases where the prior does not match the underlying population values. Reanalyzing a collection of published heritability estimates from Mousseau and Roff (1987), Kirkpatrick (2013; pers. comm.) suggested that their empirical distribution could be modelled as Beta(1.14, 1.32), corresponding to *v* = 2.46 and mode of 0.3.

We have presented two types of suitable penalties which fit well within the standard framework of REML estimation. Both achieved overall comparable reductions in loss but acted slightly differently, with penalties on correlations mainly affecting the smallest eigenvalues of the covariance matrices while penalties on canonical eigenvalues acted on both the smallest and largest values. Clearly it is the effect on the smallest eigenvalues which have the largest sampling variances which contributes most to the overall reduction in loss for a covariance matrix. An advantage of the penalty on correlations is that is readily implemented for the parameterizations commonly employed in REML estimation, and that it is straightforward to extend it to models with additional random effects and covariance matrices to be estimated or cases where traits are recorded on distinct subsets of individuals so that some residual covariances are zero. It also lends itself to scenarios where we may be less interested in a reduction in sampling variance but may want to shrink correlations towards selected target values.

Obviously, there are many other options. Mean reductions in loss obtained in a previous study, attempting to estimate tuning factors and using penalties derived assuming a Normal distribution of canonical eigenvalues or Inverse Wishart distributions of covariance or correlation matrices, again were by and large of similar magnitude (Meyer, 2011). Additional simulations in this study (not shown) using the penalty of Chung *et al*. (2015) again yielded comparable, if somewhat lower PRIAL to our penalties but required that penalties were imposed on both **Σ***_G_* and **Σ***_E_* simultaneously. In addition, like that on standard correlations such penalty was less robust, with more incidences of undesirable, negative PRIAL.

REML estimates of covariance components are biased even if no penalty is applied, as estimates are constrained to the parameter space, i.e. the smallest eigenvalues are truncated at zero or, in practice, at a small positive value to ensure estimated matrices are positive definite. As shown in Figure 3, penalization tended to increase the lower limits for the smallest canonical eigenvalues and thus also for the corresponding values of the genetic covariance matrix, thus adding to the inherent bias. Previous work examined the bias due to penalization on specific genetic parameters in more detail (Meyer and Kirkpatrick, 2010; Meyer, 2011), showing that changes from unpenalized estimates were usually well within the range of standard errors. Employing a mild penalty with fixed ESS, changes in 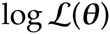 from the maximum value in an unpenalized analysis were generally small and well below significance levels (for the 90 parameters estimated in our simulation study, minus twice the change would have needed to exceed 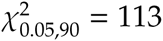 to be significant in a likelihood ratio test with an error probability of 5%). This suggests that additional bias in REML estimates due to a mild penalty on the likelihood is of minor concern and far outweighed by the benefits of a reduction in sampling variance.

Other opportunities to reduce sampling variation arise through more parsimonious modelling, e.g. by estimating **Σ***_G_* at reduced rank or assuming a factor-analytic structure. Future work should examine the scope for penalization in this context and consider the effects on model selection.

## IMPLEMENTATION

Penalized estimation for the penalties proposed for fixed values of *ν* has been implemented in our mixed model package wombat^1^ (Meyer, 2007). For 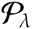 a parameterisation to the elements of the canonical decomposition is used for ease of implementation, while penalties on correlations use the standard parameterisation to elements of the Cholesky factors of the covariance matrices to be estimated. Maximization of the likelihood is carried out using the average information algorithm, combined with derivative-free search steps where necessary to ensure convergence. Example runs for a simulated data set are shown in Supplement S1.

Limited experience with applications so far has identified small to moderate effects of penalization on computational requirements compared with an unpenalized analysis using the same parameterisation, with the bulk of extra computations arising from derivative-free search steps used to check for convergence. Future work should consider an expectation-maximization algorithm for this purpose (Green, 1990). The parameterisation to the elements of the canonical decomposition, however, tended to increase the number of iterations required even without penalization. Detrimental effects on convergence behaviour when parameterising to eigenvalues of covariance matrices have been reported previously (Pinheiro and Bates, 1996).

Convergence rates of iterative maximum likelihood analyses are dictated by the shape of the likelihood function. Newton-Raphson type algorithms, including the average information algorithm, involve a quadratic approximation of the likelihood. When this is not the appropriate shape, the algorithm may become ‘stuck’ and fail to locate the maximum. This happens quite frequently for (unpenalized) multivariate analyses comprising more than a few traits when covariance matrices have eigenvalues close to zero, i.e. estimates are at the boundary of the parameter space. For such cases, additional maximisation steps using alternative schemes, such as expectation maximization type algorithms or a derivative-free search, are usually beneficial. For small data sets, we expect the likelihood surface around the maximum to be relatively flat. Adding additional ‘information’ through the assumed prior distribution (*a.k.a*. the penalty) then can improve convergence by adding curvature to the surface and creating a more distinct maximum. Conversely, too stringent a penalty may alter the shape of the surface sufficiently so that a quadratic approximation may not be successful. Careful checking of convergence should be an integral part of any multivariate analysis, penalized or not.

## CONCLUSIONS

We propose a simple but effective modification of standard multivariate maximum likelihood analyses to ‘improve’ estimates of genetic parameters: Imposing a penalty on the likelihood designed to reduce sampling variation will yield estimates that are on average closer to the population values than unpenalized values. There are numerous choices for such penalties. We demonstrate that those derived under the assumption of a Beta distribution for scale-free function of the covariance components to be estimated, namely generalized heritabilities (*a.k.a* canonical eigenvalues) and genetic correlations, are well suited and tend not to distort estimates of the total, phenotypic variance. In addition, this allows the stringency of penalization to be regulated by a single parameter, known as effective sample size of the prior in a Bayesian context. Aiming at moderate rather than optimal improvements in estimates, suitable default values for this parameters can be identified which yield a mild penalty. This allows us to abandon the laborious quest to identify tuning factors suited to particular analyses. Choosing the penalty to be sufficiently mild can all but eliminate the risk of detrimental effects, and results in only minor changes in the likelihood, compared to unpenalized analyses. Mildly penalized estimation is recommended for multivariate analyses in quantitative genetics considering more than a few traits to alleviate the inherent effects of sampling variation.

## APPENDIX

### Derivatives of partial auto-correlations

Partial auto-correlations (see (3)) can be written as

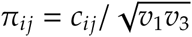

with

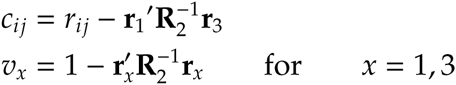

This gives partial derivatives of *π_ij_* with respect to parameters *θ_k_* and *θ_m_*

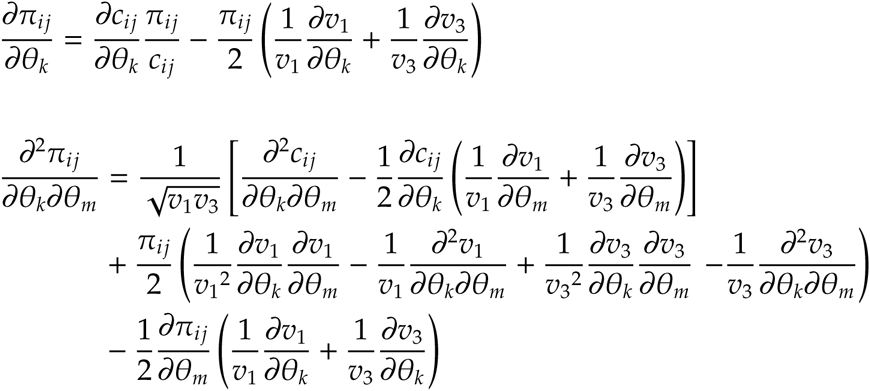

Derivatives of the components are

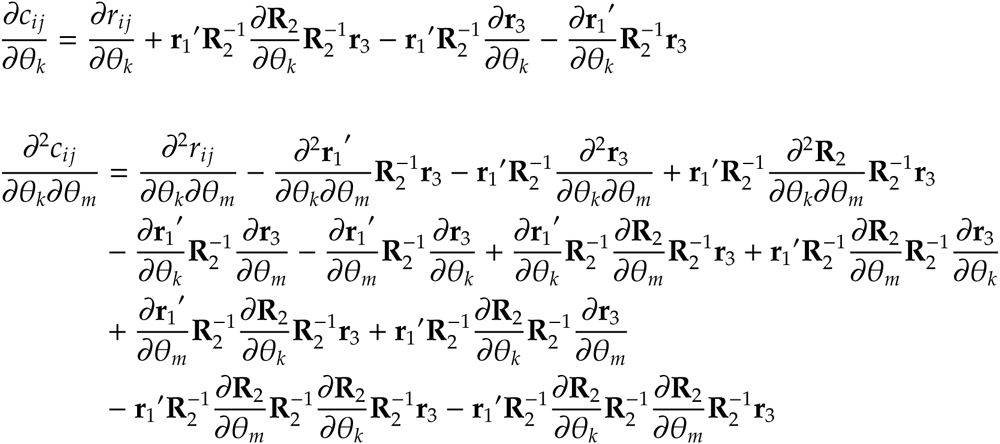

and

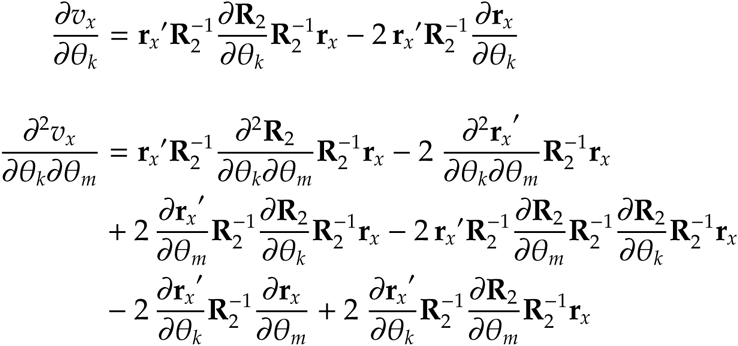

Decompose the correlation matrix as **R** = **S**^−1^**ΣS**^−1^ with **S** = Diag{*s_i_*} the diagonal matrix of standard deviations for covariance matrix **Σ**. This gives the required derivatives of the correlation matrix

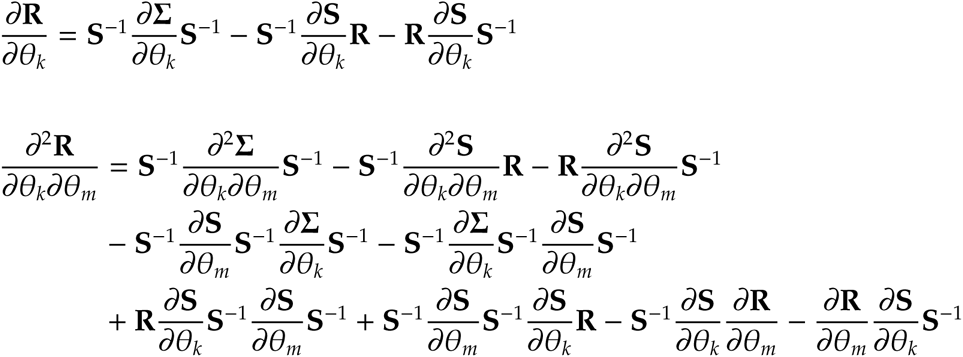

Finally, assuming derivatives of variances *σ_ii_* are available, the required derivatives of standard deviations 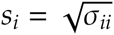 are

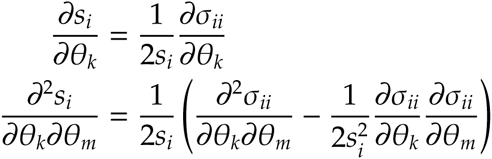

When estimating elements of **Σ** directly, only ∂*σ_ii_*/∂*σ_ii_* = 1 are nonzero. Derivatives of covariance matrices when employing a parameterisation to the elements of their Cholesky factors are given by Meyer and Smith (1996).

### Population values

Population values for the 13 sets of heritabilities used are summarized in Table 3. The six constellations of genetic (*r_Gij_*) and residual (*r_Eij_*) correlations between traits *i* and *j* (*i* ≠ *j*) were obtained as:

i. *r_Gij_* = *r_Eij_* = 0,
ii. *r_Gij_* = 0.5 and *r_Eij_* = 0.3,
iii. *r_Gij_* = 0.7^|^*^i^*^−^*^j^*^|^ and *r_Eij_* = 0.5 + 0.05*i* (−1)*^j^*,
iv. *r_Gij_* = −0.7^|^*^i^*^−^*^j^*^|^ + 0.02*i* and *r_Eij_* = 0.5 + (−0.2)^|^*^i^*^−^*^j^*^|^,
v. *r_Gij_* = *r_Eij_* = 0.7 for *i*, *j* ∈ [3, 7] and *r_Gij_* = *r_Eij_* = 0.3 otherwise, and
vi. *r_Gij_* = *r_Eij_* = 0.6 for |*i* − *j*| = 1, and *r_Gij_* = *r_Eij_* computed from (4) with *π_ij_* = 0.4 otherwise.

**Table 3:** Population values (×100) for sets of heritabilities

| Set | Trait number |  |  |  |  |  |  |  |  |
| --- | --- | --- | --- | --- | --- | --- | --- | --- | --- |
|  | 1 | 2 | 3 | 4 | 5 | 6 | 7 | 8 | 9 |
| A | 40 | 40 | 40 | 40 | 40 | 40 | 40 | 40 | 40 |
| B | 60 | 55 | 50 | 45 | 40 | 35 | 30 | 25 | 20 |
| C | 90 | 60 | 50 | 50 | 30 | 30 | 20 | 20 | 10 |
| D | 75 | 70 | 60 | 50 | 40 | 30 | 20 | 10 | 5 |
| E | 70 | 70 | 70 | 40 | 40 | 40 | 10 | 10 | 10 |
| F | 20 | 20 | 20 | 20 | 20 | 20 | 20 | 20 | 20 |
| G | 35 | 30 | 25 | 20 | 20 | 20 | 15 | 10 | 5 |
| H | 60 | 50 | 10 | 10 | 10 | 10 | 10 | 10 | 10 |
| I | 50 | 50 | 20 | 15 | 15 | 10 | 10 | 5 | 5 |
| J | 80 | 40 | 10 | 10 | 10 | 10 | 10 | 5 | 5 |
| K | 30 | 30 | 25 | 25 | 20 | 15 | 15 | 10 | 10 |
| L | 35 | 30 | 30 | 20 | 20 | 15 | 15 | 15 | 10 |
| M | 10 | 10 | 10 | 30 | 30 | 30 | 50 | 50 | 50 |

Phenotypic variances were equal to 1 throughout for I) and set to 2, 1, 3, 2, 1, 2, 3, 1 and 2 for traits 1 to 9 otherwise.

## Footnotes

1 available from http://didgeridoo.une.edu.au/km/wombat.php

## LITERATURE CITED

Anderson, T. W., 1984 An Introduction to Multivariate Statistical Analysis. Wiley, New York, 2nd edition.

Barnard, J., R. McCulloch, and X. Meng, 2000 Modeling covariance matrices in terms of standard deviations and correlations, with applications to shrinkage. Stat. Sin. 10: 1281–1312.

Bickel, P. J., and E. Levina, 2008 Regularized estimation of large covariance matrices. Ann. Stat. 36: 199–227.

Bickel, P. J., and B. Li, 2006 Regularization in statistics. Test 15: 271–303.

Bouriga, M., and O. Féron, 2013 Estimation of covariance matrices based on hierarchical inverse-Wishart priors. J. Statist. Plann. Inference 143: 795–808.

Cheverud, J. M., 1988 A comparison of genetic and phenotypic correlations. Evolution 42: 958–968.

Chung, Y., A. Gelman, S. Rabe-Hesketh, J. Liu, and V. Dorie, 2015 Weakly informative prior for point estimation of covariance matrices in hierarchical models. Journal of Educational and Behavioral Statistics 40: 136–157.

Daniels, M. J., and R. E. Kass, 2001 Shrinkage estimators for covariance matrices. Biometrics 57: 1173–1184.

Daniels, M. J., and M. Pourahmadi, 2009 Modeling covariance matrices via partial auto-correlations. J. Multiv. Anal. 100: 2352–2363.

Deng, X., and K.-W. Tsui, 2013 Penalized covariance matrix estimation using a matrix-logarithm transformation. J. Comp. Graph. Stat. 22: 494–512.

Fisher, T. J., and X. Sun, 2011 Improved Stein-type shrinkage estimators for the high-dimensional multivariate normal covariance matrix. Comp. Stat. Dat. Anal. 55: 1909–1918.

Gaskins, J. T., M. J. Daniels, and B. H. Marcus, 2014 Sparsity inducing prior distributions for correlation matrices of longitudinal data. J. Comp. Graph. Stat. 23: 966–984.

Gelman, A., 2006 Prior distributions for variance parameters in hierarchical models. Bayesian Analysis 1: 515–533.

Gilmour, A. R., R. Thompson, and B. R. Cullis, 1995 Average Information REML, an efficient algorithm for variance parameter estimation in linear mixed models. Biometrics 51: 1440–1450.

Green, P. J., 1987 Penalized likelihood for general semi-parametric regression models. Int. Stat. Rev. 55: 245–259.

Green, P. J., 1990 On use of the EM for penalized likelihood estimation. J. Roy. Stat. Soc. B 52: 443–452.

Hayes, J.F., and W.G. Hill, 1980 A reparameterisation of a genetic index to locate its sampling properties. Biometrics 36: 237–248.

Hayes, J. F., and W. G. Hill, 1981 Modifications of estimates of parameters in the construction of genetic selection indices (‘bending’). Biometrics 37: 483–493.

Hill, W. G., 2010 Understanding and using quantitative genetic variation. Phil. Trans. Roy. Soc. B 365: 73–85.

Hsu, C. W., M. S. Sinay, and J. S. J. Hsu, 2012 Bayesian estimation of a covariance matrix with flexible prior specification. Ann. Inst. Statist. Math. 64: 319–342.

Huang, A., and M. P. Wand, 2013 Simple marginally noninformative prior distributions for covariance matrices. Bayesian Analysis 8: 439–452.

Huang, J.Z., N. Liu, M. Pourahmadi, and L. Liu, 2006 Covariance matrix selection and estimation via penalised normal likelihood. Biometrika 93: 85–98.

James, W., and C. Stein, 1961 Estimation with quadratic loss. In J. Neiman, editor, Proceedings of the Fourth Berkeley Symposium on Mathematical Statistics and Probability. University of California Press, 361–379.

Joe, H., 2006 Generating random correlation matrices based on partial correlations. J. Multiv. Anal. 97: 2177–2189.

Johnson, N., S. Kotz, and N. Balakrishnan, 1995 Continuous Univariate Distributions, volume 2 of Wiley Series in Probability and Mathematical Statistics: Applied Probability and Statistics Section. Wiley & Sons. Inc, New York, 2nd edition.

Koots, K. R., J. P. Gibson, and J. W. Wilton, 1994 Analyses of published genetic parameter estimates for beef production traits. 2. Phenotypic and genetic correlations. Anim. Breed. Abstr. 62: 825–853.

Lawley, D. N., 1956 Tests of significance for the latent roots of covariance and correlation matrices. Biometrika 43: 128–136.

Ledoit, O., and M. Wolf, 2004 A well-conditioned estimator for large-dimensional covariance matrices. J. Multiv. Anal. 88: 365–411.

Ledoit, O., and M. Wolf, 2012 Nonlinear shrinkage estimation of large-dimensional covariance matrices. The Annals of Statistics 40: 1024–1060.

Lin, S. P., and M. D. Perlman, 1985 A Monte Carlo comparison of four estimators of a covariance matrix. In P. R. Krishnaish, editor, Multivariate Analysis, volume 6. North-Holland, Amsterdam, 411–428.

Meyer, K., 2007 WOMBAT—a tool for mixed model analyses in quantitative genetics by REML. J. Zhejiang Univ. SCIENCE B 8: 815–821.

Meyer, K., 2011 Performance of penalized maximum likelihood in estimation of genetic covariances matrices. Genet. Sel. Evol. 43: 39.

Meyer, K., and M. Kirkpatrick, 2010 Better estimates of genetic covariance matrices by ‘bending’ using penalized maximum likelihood. Genetics 185: 1097–1110.

Meyer, K., M. Kirkpatrick, and D. Gianola, 2011 Penalized maximum likelihood estimates of genetic covariance matrices with shrinkage towards phenotypic dispersion. Proc. Ass. Advan. Anim. Breed. Genet. 19: 87–90.

Meyer, K., and S. P. Smith, 1996 Restricted maximum likelihood estimation for animal models using derivatives of the likelihood. Genet. Sel. Evol. 28: 23–49.

Morita, S., P. F. Thall, and P. Müller, 2008 Determining the effective sample size of a parametric prior. Biometrics 64: 595–602.

Mousseau, T.A., and D.A. Roff, 1987 Natural selection and the heritability of fitness components. Heredity 59: 181–197.

Pinheiro, J.C., and D.M. Bates, 1996 Unconstrained parameterizations for variance-covariance matrices. Stat. Comp. 6: 289–296.

Powell, M., 2006 The NEWUOA software for unconstrained optimization without derivatives. In G. Pillo and M. Roma, editors, Large-Scale Nonlinear Optimization, volume 83 of *Nonconvex Optimization and Its Applications*. 255–297.

Rapisarda, F., D. Brigo, and F. Mercurio, 2007 Parameterizing correlations: a geometric interpretation. IMA J. Managem. Math. 18: 55.

Roff, D. A., 1995 The estimation of genetic correlations from phenotypic correlations - a test of Cheveruds conjecture. Heredity 74: 481–490.

Rothman, A. J., E. Levina, and J. Zhu, 2010 A new approach to Cholesky-based covariance regularization in high dimensions. Biometrika 97: 539–550.

Schäfer, J., and K. Strimmer, 2005 A shrinkage approach to large-scale covariance matrix estimation and implications for functional genomics. Stat. Appl. Genet. Mol. Biol. 4: 32.

Stein, C., 1975 Estimation of a covariance matrix. In Reitz lecture of the 39th Annual Meeting of the Institute of Mathematical Statistics. Atlanta.

Thompson, R., S. Brotherstone, and I. M. S. White, 2005 Estimation of quantitative genetic parameters. Phil. Trans. Roy. Soc. B 360: 1469–1477.

Waitt, D. E., and D. A. Levin, 1998 Genetic and phenotypic correlations in plants: a botanical test of Cheverud’s conjecture. Heredity 80: 310–319.

Warton, D. I., 2008 Penalized normal likelihood and ridge regularization of correlation and covariance matrices. J. Amer. Stat. Ass. 103: 340–349.

Witten, D. M., and R. Tibshirani, 2009 Covariance-regularized regression and classification for high dimensional problems. J. Roy. Stat. Soc. B 71: 615–636.

Won, J.-H., J. Lim, S.-J. Kim, and B. Rajaratnam, 2013 Condition-number-regularized covariance estimation. J. Roy. Stat. Soc. B 75: 427–450.

Ye, R. D., and S. G. Wang, 2009 Improved estimation of the covariance matrix under Stein’s loss. Stat. Prob. Lett. 79: 715–721.

Zhang, X., W. J. Boscardin, and T. R. Belin, 2006 Sampling correlation matrices in Bayesian models with correlated latent variables. J. Comp. Graph. Stat. 15: 880–896.

